# Design of high specificity binders for peptide-MHC-I complexes

**DOI:** 10.1101/2024.11.28.625793

**Authors:** Bingxu Liu, Nathan F. Greenwood, Julia E. Bonzanini, Amir Motmaen, Jazmin Sharp, Chunyu Wang, Gian Marco Visani, Dionne K. Vafeados, Nicole Roullier, Armita Nourmohammad, K. Christopher Garcia, David Baker

## Abstract

Class I MHC molecules present peptides derived from intracellular antigens on the cell surface for immune surveillance, and specific targeting of these peptide-MHC (pMHC) complexes could have considerable utility for treating diseases. Such targeting is challenging as it requires readout of the few outward facing peptide antigen residues and the avoidance of extensive contacts with the MHC carrier which is present on almost all cells. Here we describe the use of deep learning-based protein design tools to *de novo* design small proteins that arc above the peptide binding groove of pMHC complexes and make extensive contacts with the peptide. We identify specific binders for ten target pMHCs which when displayed on yeast bind the on-target pMHC tetramer but not closely related peptides. For five targets, incorporation of designs into chimeric antigen receptors leads to T-cell activation by the cognate pMHC complexes well above the background from complexes with peptides derived from proteome. Our approach can generate high specificity binders starting from either experimental or predicted structures of the target pMHC complexes, and should be widely useful for both protein and cell based pMHC targeting.

## Introduction

MHC-I molecules are cell-surface proteins that present peptides derived from intracellular proteins. The recognition of peptides displayed on MHC-I by the T-cell receptor enables the immune system to detect foreign proteins within cells and destroy tumor and viral infected cells. From the therapeutic perspective, targeting pMHCs with engineered cells (*1*, *2*) or proteins–for example, Bispecific T cell Engagers (BiTEs) (*3*); (*4*)–is attractive as it provides a unique opportunity to distinguish cells based on their intracellular proteins. T-cell receptors can be used for such targeting, but TCRs with the necessary specificity have not been identified for many targets of therapeutic interest, engineering TCRs with new specificities has been very challenging, and for protein based therapeutic strategies, TCR extracellular domains are quite difficult to produce in soluble form. The diversity of MHC alleles and antigenic peptides across patient populations necessitates the development of hundreds to thousands of effective and specific binders to achieve broad patient coverage. Current methods rely on empirical TCR screening from patient samples (*5*) or single-chain variable fragment (scFv) libraries (*1*, *3*) are costly, labor-intensive, and time-consuming, making them economically infeasible for covering all pMHC targets, especially for smaller patient subpopulations. Methods for rapidly generating small stable proteins that recognize specific pMHCs of interest could have considerable therapeutic utility as the recognition domains in both chimeric antigen receptors (CARs) for cell based therapies, and in protein based therapies such as BiTEs.

We reasoned that *de novo* protein design could provide a powerful approach to the specific pMHC recognition problem. Recent advances in deep learning-driven binder design have enabled the structure based design of high affinity and specificity binders against a wide range of folded (*6–8*) and disordered (*9*, *10*) protein targets. These designed binders are small (<150 amino acids) and very stable proteins which can be readily expressed in *E. coli*, and the binding modes of computationally designed binders are generally very close to experimentally determined structures of the complexes, enabling inference of binding specificity and further optimization without need for time consuming and expensive experimental structure determination. We reasoned that if such binders could be generated against pMHCs there could be multiple advantages in stability, engineerability, and manufacturability, and set out to explore the computational design of high affinity and specificity pMHC binders.

## Results

Critical to pMHC recognition for therapeutic applications is for binding to be dependent on the identity of the peptide, as MHCs can present other peptides from the proteome that share similar sequences (**Fig. 1a**); indeed, high specificity TCRs and scFvs make extensive interactions with the presented peptides (*3*, *5*). To specifically target the peptide in pMHC complexes, we used the generative AI protein design method RFdiffusion to generate protein backbones specifying the upward facing (out of the MHC groove) residues of the peptide as hotspots (**Fig.1b**). Starting from random Gaussian distributions of amino acid residues placed adjacent to the peptide, diffusive denoising trajectories generated a wide variety of small protein backbones that arc above the central peptide binding groove of the MHC (**supp Fig.1a**). From tens of thousands of independent trajectories, we selected backbones capable of hosting side chains making many contacts with the peptide based on contact molecular surface (CMS) (*11*) (**supp Fig.1b**). Following sequence design with ProteinMPNN, we selected for further consideration the designs which AlphaFold2 (AF2) (*12*) predicted to fold and bind as designed. Once such designs were identified for the first few targets, we explored using partial diffusion (*13*) to adapt the most promising scaffolds to new pMHC targets (**Fig. 1c**). Starting from a shape complementary binder to one member of a protein family, partial diffusion can rapidly generate high specificity binders to other family members (*6*); this partial diffusion based customization of promising scaffolds for new targets could reduce computational cost as only a small fraction of *de novo* RFdiffusion trajectories generate backbones which make extensive contacts with the peptide while largely avoiding the MHC.

**Figure 1:**
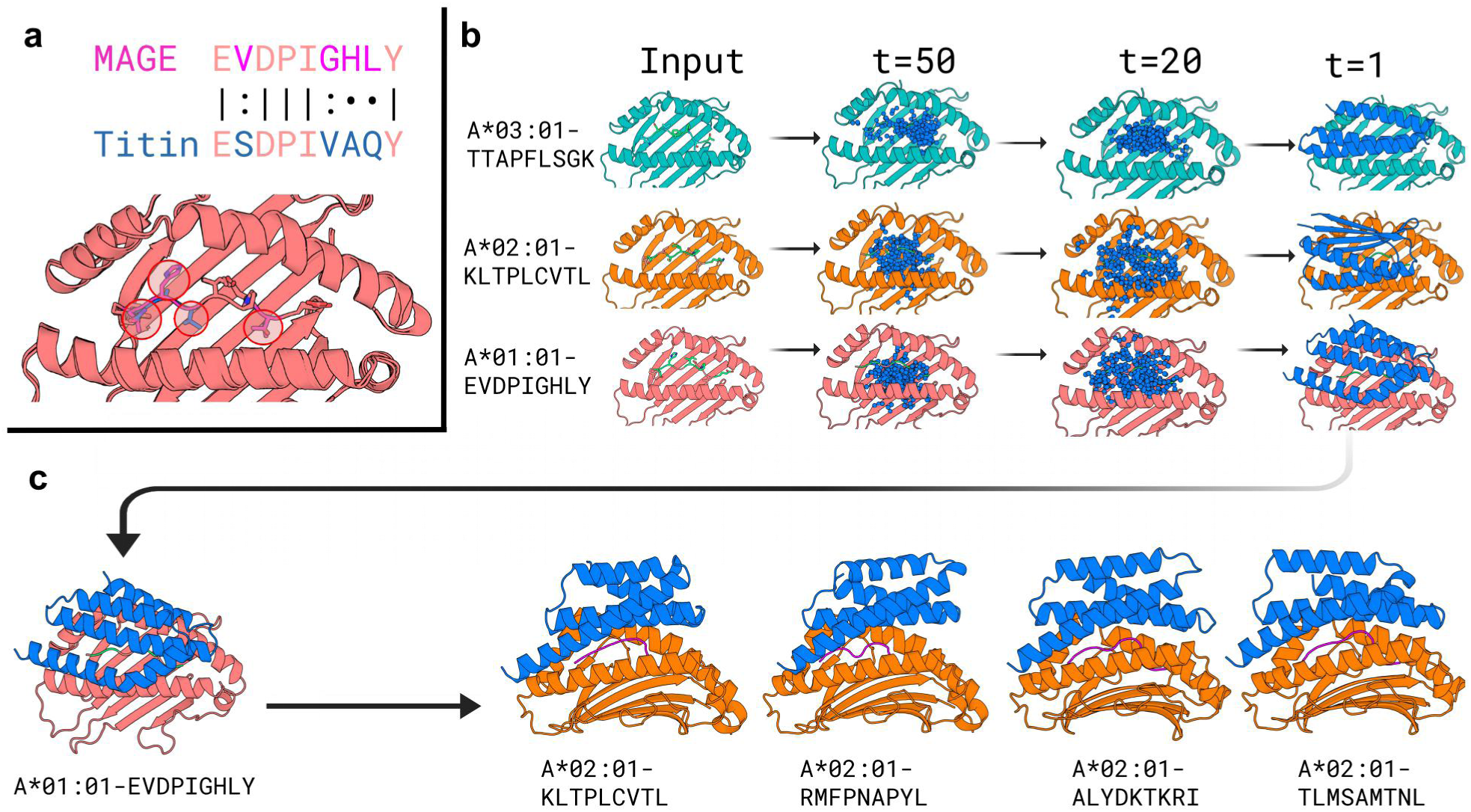
Diffusion of pMHC binders. **a**, pMHC structure and design challenge. The goal is to distinguish a target peptide (in this example, MAGE) from a closely related off-target (Titin). The positions that differ between the two peptides are circled in the pMHC structure model at the bottom. **b,** Representative diffusion trajectories and design models for three different pMHC targets. Left two columns, target pMHc identities and structures. Columns 3 and 4, intermediate steps in diffusion denoising trajectories starting from completely random residue distributions. Right column, fully denoised design model backbones. **c,** Partial diffusion of design scaffolds with desirable properties (interacting over the full length of the peptide, but making few interactions with MHC) to more efficiently generate binders to related targets.

For pMHC recognition for therapeutic applications, specificity is of the utmost importance. We used two computational approaches to evaluate the specificity of the RFdiffusion/ProteinMPNN generated binders for the intended target peptide. First, we used ProteinMPNN to evaluate the effect of mutating each amino acid of the target peptide for each designed protein-pMHC complex. We selected designs for which the peptide amino acid had high probability at each position, and for which substitutions at each position reduced this probability considerably (**supp Fig.1c**). Second, for each target, we identified closely-related peptides (see methods) in the human proteome that are presented by the target HLA allele. We used a finetuned AF2 model to predict the structure of each design with both the targeted and off-target peptides in complex with the MHC, and selected designs predicted to bind the on-target peptide (lower pAE scores) considerably more confidently than the off-target peptides (higher pAE scores) (**supp Fig.1d**).

To evaluate the ability of this design pipeline to generate binders to a wide range of pMHC complexes, we selected a structurally diverse set of ten complexes comprising HLA alleles A*01:01, A*02:01 and A*03:01 presenting 9– and 10-mer peptides from viral proteins, tumor associated proteins, and neoantigens. For each target, we obtained oligonucleotide pools encoding 200-12,000 designs, displayed the designs on yeast, and selected those that specifically recognized the targeted pMHC, but not 2-4 closely related peptides (**Supp Table.1**) on the same MHC (referred to off-target peptides below), by dual color fluorescent cell sorting (FACS) (**Supp Fig.2a**). Top designs were selected through NGS enrichment analysis or clonal selection (**Fig.2a**). For seven of the ten pMHC complexes, we used *de novo* diffusion, and identified designs with a range of topologies (**Fig.2a, left**) and peptide binding interfaces (**Fig.2a, right**) that specifically bound the target peptide, with reduced binding to off-target peptides on the same HLA (**Fig.2b**). For three of the targets, we used the partial diffusion approach starting from one of the *de novo* designs (**Fig.2c top**); we again identified specific on-target binders in each case (**Fig.2c down**). While we were not able to solve structures experimentally of the designs in complex with pMHC, AF3 and Chai-1 (*14*, *15*) predictions of the structures of the designed proteins in complex with pMHC were very similar to the design models (**Supp Table 1, Supp Fig.2b and c, right;** for simplicity we refer to these interchangeably as the design model below). We describe the results for each class of targets in the following paragraphs.

To evaluate the ability of the design pipeline to generate binders to viral peptide-HLA complexes, we chose three previously characterized epitopes from different pathogens presented on A*02:01: SARS-CoV1/GLMWLSYFV (*16*), YFV/LLWNGPIAV (*17*), HIV/KLTPLCVTL (*18*). For all three targets, we identified designs that distinguish the target peptides from closely related off-target peptides (**Fig.2b**). To assess whether the designs can maintain the specificity as the binding domain for chimeric antigen receptors (CARs), we made CAR constructs (*19*) and expressed them on the surface of Jurkat cells. While not all of the CARs expressed, top specific hits that expressed have specific pMHC binding profiles (**Fig.2b**). In the design models (**Fig.2a**), the binders make extensive contacts with the target peptide (**Supp Table 2**). As with TCR-pMHC complexes, target sequence specificity arises in two ways: first by direct interactions between designed side chains and target peptide side chains, and second, by specific recognition of the conformation of the bound peptide by hydrogen bonds with the target peptide backbone, which is also determined by the peptide amino acid sequence. For example, the SARS design buries the target peptide W4, L5 and Y7 in a hydrophobic pocket, and makes flanking hydrogen bonds with the peptide backbones; the YFV design makes hydrogen bonds with N4 and hydrophobic interactions with I7 of the peptide; and the HIV design makes hydrogen bonds with K1, T8, and the peptide backbone. We next evaluated the ability of our design pipeline to target diverse tumor associated antigens (TAAs): WT1 (RMFPNAPYL/A*02:01) (*20*, *21*), PAP (TLMSAMTNL/A*02:01) (*22*), and the neoantigen CTNNb1 (TTAPFLSGK/A*03:01 with S5F mutation) (*23*). We identified specific designs for all three peptides (**Fig.2b)**; as found for the viral antigens, the design models show extensive interaction between peptides and designed binders (**Fig.2a)**. The CTNNb1 design makes extensive contact with the mutated Phe residue, through cation-pi, pi-pi, and other hydrophobic interactions (**Fig.2a)**, likely contributing to the specific binding observed for the S5F peptide but not the Wild-Type (WT) CTNNb1 peptide (**Fig.2b)**.

To test the partial diffusion approach, we started from a design that binds the MAGE-A3 peptide (A*01:01-EVDPIGHLY) (*24*, *25*) but not the off-target Titin peptide (A*01:01-ESDPIVAQY) (*26*) (**Fig.2c, top**). Starting from this scaffold, we generated specific binders for other 9– and 10-residue tumor associated antigens presented by A*02:01: the gp100 peptide YLEPGPVTA (*27*) and the Mart-1 peptide (A2L) ELAGIGILTV (*28*). As only a small fraction of potential pMHC targets have experimentally determined structures, we also explored the possibility of using predicted pMHC structures as targets for binder design. The PRAME protein is highly expressed in multiple types of tumors, and a peptide derived from PRAME (ALYVDSLFFL) (*29*) is displayed on HLA-A*02:01. Despite the strong therapeutic relevance and wide patient population coverage, no high-resolution structure has been determined. AF3 predictions of the complex are confident but display some flexibility in the central region of the peptide, consistent with the intrinsic flexibility of 10-mer peptides in the MHC. For design, we experimented with using 5 AF3 predicted structures as starting points. For all three peptide-MHC targets (gp100, Mart-1, and PRAME), we identified specific binders that make extensive and diverse hydrophobic and hydrogen bonding interactions with the target peptide (**Fig.2c**).

To evaluate the behavior of our designs as soluble proteins, we expressed the HIV and MAGE-A3 binder designs shown in Fig.2 in *E. coli* and purified them using nickel-NTA chromatography (**Supp Fig.2b and c**). The purified designs eluted as single peaks in size exclusion chromatography around expected size. We measured the binding of the designs to their cognate pMHCs using surface plasmon resonance experiments and found the binding affinities to be in the single to double digit nanomolar range (**Supp Fig.2b and c**).

**Figure 2:**
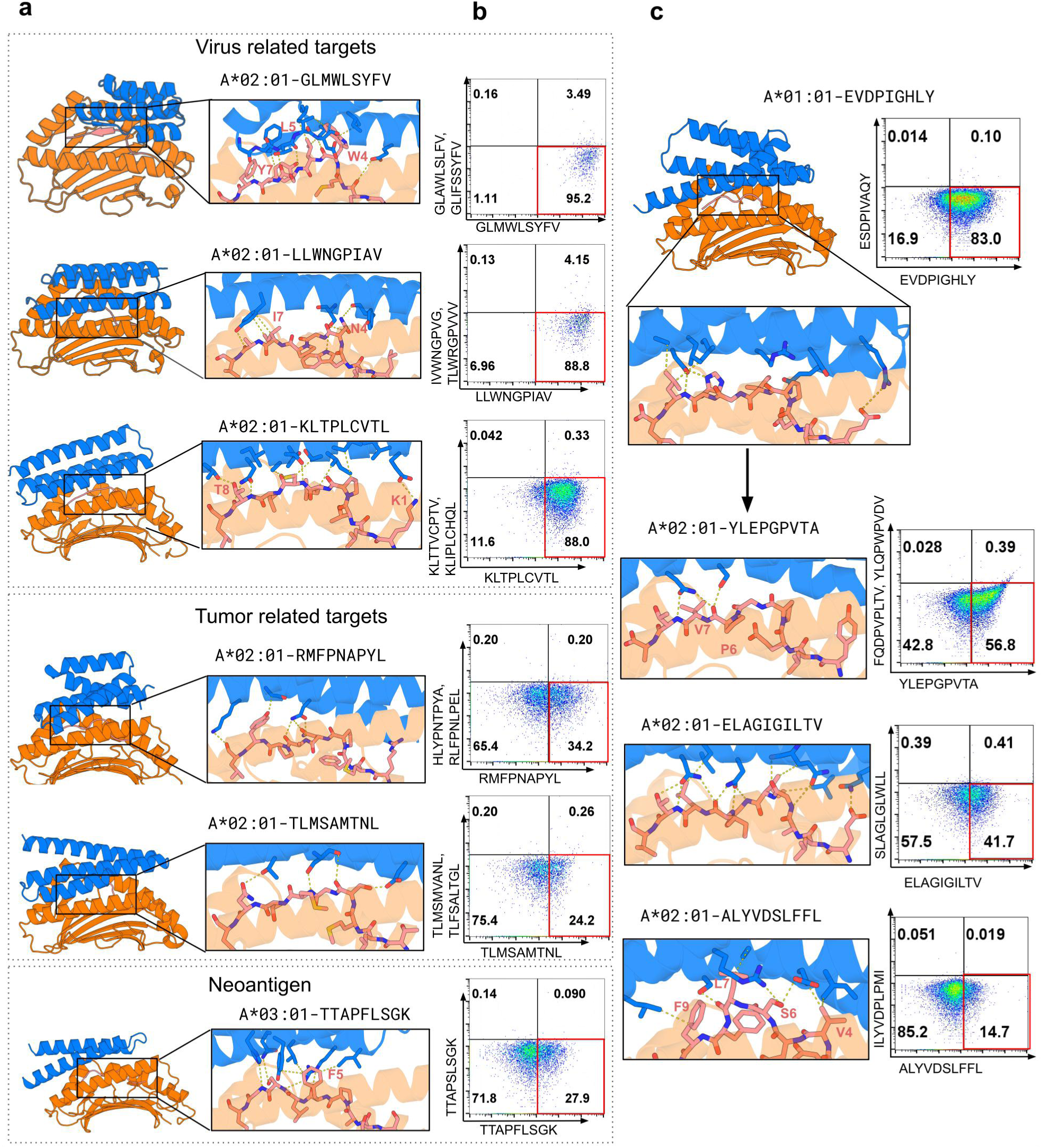
Design models and binding specificity. **a**, Design models. Left, overall structure; right, zoom in on peptide binding region. pMHC in orange, peptide in sticks, and binder in blue. HLA allele and peptide sequence are specified above the zoom-in view. **b,** Flow cytometry of cells displaying the design incubated with on-target pMHC tetramer (x-axis) and 1 or 2 off-target tetramers (y-axis) at 10nM concentration. Staining in the lower right quadrant indicates specific on-target binding. Rows 1 and 2, individual designs displayed on yeast; rows 3-6, CARs incorporating designs on Jurkat cells. **c,** Partial diffusion of design at top to targets below (left panels) and corresponding Jurkat straining (right).

We next evaluated the ability of the designs to enable specific T-cell activation by the target pMHC when incorporated into chimeric antigen receptors. To be effective and safe in cell therapy settings, such CARs incorporating designed proteins must mediate specific activation by the target peptide loaded on the target HLA, but not by the thousands of other peptides from the proteome loaded on the same HLA, or by peptides on different HLAs. To evaluate such activation specificity, we incorporated the binders into CARs and expressed them on Jurkat cells, incubating them with 293T cells that were treated with on-target or off-target peptides. 293T cells have an intact antigen presentation system, and hence present a wide range of peptides derived from self proteins on their surfaces; requiring selective activation by pulsing peptides in this system is thus more stringent than using cells that cannot present self peptides due to defects in presentation.

Among the designed MAGE binders, the MAGE-513 design resulted in strong and specific activation only by MAGE-A3 peptide stimulation but not by the off-target Titin peptide, or peptides derived from the intracellular proteome on HLA-A*01:01 (**Fig.3a**). Despite binding to target tetramer specifically (**Supp Fig.3a**), Other designs had either weak signaling (**Supp Fig.3b,** MAGE-282) or background activation (**Supp Fig.3b,** hit-4) likely due to responses to other peptides from the proteome loaded on the same HLA as only HLA-A*01:01 expressing 293T cells give the background activation. In the design model (**Fig.3b)**, MAGE-513 engages the MAGE peptide through hydrogen bonds with the side chain of H7 and backbone of L8 (**Fig.3b),** and through hydrophobic interactions (mediated by design L37 and L86) with the sidechain of peptide L8 (there are also hydrogen bonds between R33 of the design and N66 of the HLA) **(Fig.3b)**. D29 of the design is close to P4 of the peptide without making a clear interaction (**Fig.3b)**; this residue likely plays a gatekeeper role that contributes to binding specificity by clashing with off target peptides that contain bulky residues at site 4. Consistent with the design model and predicted structure, alanine mutations of the most extensively interacting residues on the peptide (I5, H7, L8) disrupt signaling (**Fig.3c)**. Similarly, CARs with alanine mutations in designed binder residues (L37, D40, L86) which interact most closely with the peptides show reduced activation compared to WT MAGE-513 upon stimulation (**Fig.3d)**. Mutation of D29 does not influence signal upon pulsing with MAGE peptide but does have higher background signal when co-incubated with HLA-A*01:01 expressing 293T cells likely due to increased cross-reactivity, consistent with a gate-keeper role (**Fig.3d**).

**Figure 3:**
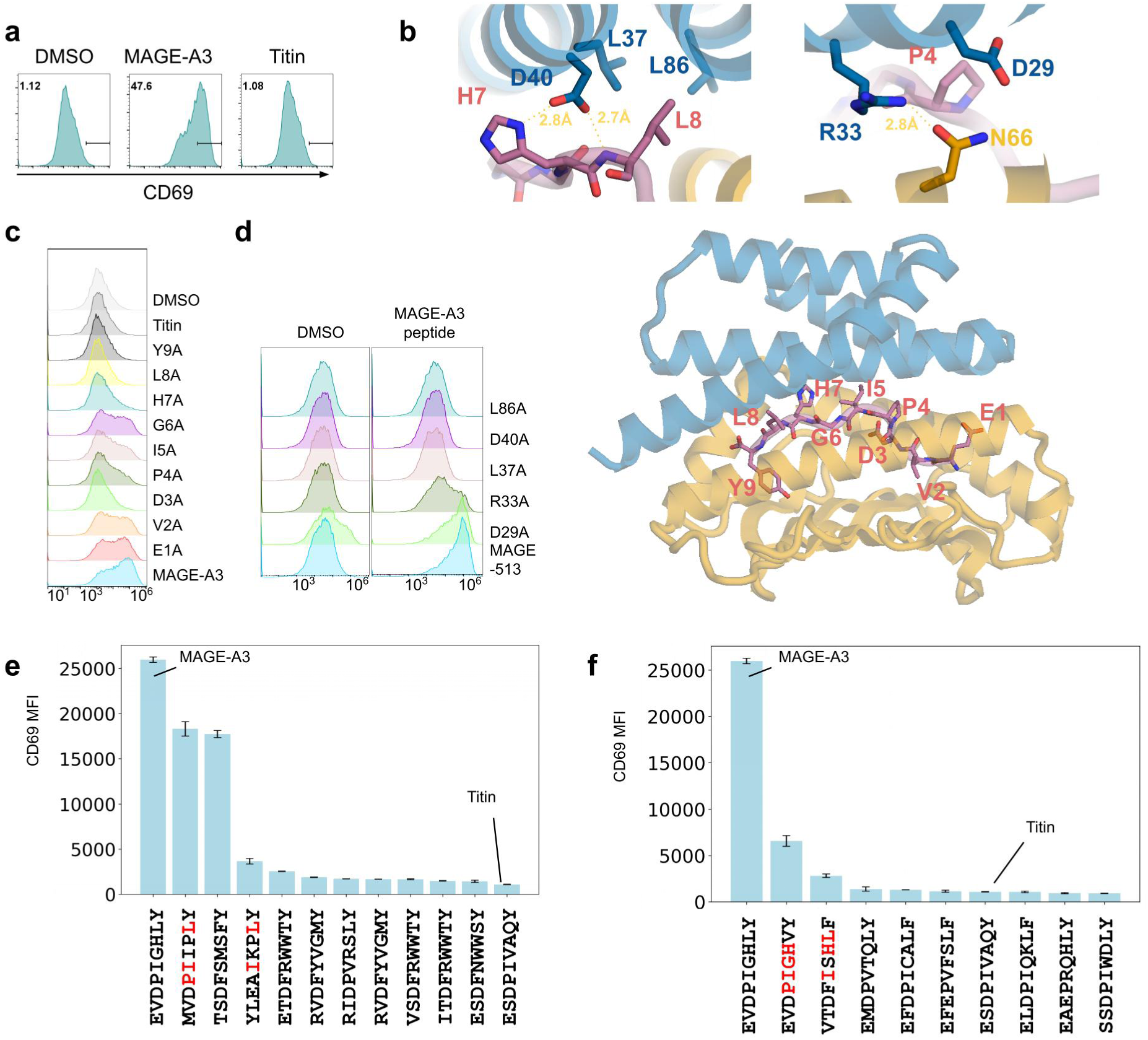
Selective activation of T-cells expressing designed CARs targeting MAGE-A3. Activation of Jurkat cells expressing the MAGE-513 CAR by 293T cells expressing HLA-A*01:01 pulsed with 5uM of different peptides measured through CD69 expression level. **a,** Histograms of CD69 expression levels following treatment with 5uM MAGE, the closely related Titin peptide or DMSO; the MAGE peptide leads to considerable activation while the Titin peptide is similar to the DMSO control. Horizontal black bars represent CD69 positive population; the fraction of cells within this range is indicated at top left of each panel. **b,** Design models of the MAGE-513/MAGE peptide/HLA-A*01:01 complex **(lower panel)** and zoom-in view of the key residues mediating the interaction **(upper panels). c,** Histograms of CD69 expression levels following pulsing with MAGE-A3 single alanine mutants (D3A indicates mutation of the Asp at position 3 in the peptide to alanine) or DMSO. d,Histograms of CD69 expression levels following pulsing MAGE-513 variant CARs with MAGE-A3 peptide or DMSO. **e and f,** CD69 levels of Jurkat cells expressing MAGE-513 CAR upon pulsing with top ranked peptides from yeast binding screen **(e)** or from sequence similarity search **(f).** Identity to the MAGE peptide is indicated in red.

Comprehensive understanding of the pMHC binder activation landscape requires library-wide scanning of diverse peptides (*30*). To further characterize the specificity of the MAGE-A3 design, we used a comprehensive library of peptide-HLA-A*01:01 complexes displayed on yeast. Two of the top three activating peptides found probing this library with the MAGE-A3 design, and then measuring activation of MAGE-A3 CAR T cells by the identified peptides, have similar outward facing sidechains as the MAGE-A3 peptide (**Fig.3e, shown in red**). Thus, similar to TCRs, peptides that cross-react and activate signaling through our *de novo* binders have similar sequences, indicating that our design pipeline can generate specificity for a very small subset of possible sequences within a predictable range. Following up on this observation, we carried out a second scan of cross-activating peptides in the human proteome based on sequence similarity (*31*). The most activating peptides (EVDPIGHVY and VTDFISHLF) again are among the most similar sharing outward facing residues I5, H7, L8 in sequence (**Fig.3f**), consistent with the alanine scanning results.

Given these promising results with our designs against the MAGE pMHC, we explored activation of signaling by cells expressing CARs incorporating designs for additional targets. We observed selective activation of signaling for CARs containing designs against gp100, WT1, MART-1, and HIV antigens (**Fig.4a-e, Supp Fig.4a,** compare “Target peptide” to “DMSO” histograms). For WT1, gp100, and MART-1, pulsing 293T cells with the targeted peptide activated at the same or higher levels than pulsing with almost all single alanine peptide variants (**Fig4.a-e;** for HIV an alanine mutant increased activation (**Supp Fig.4e**), with activation patterns consistent with the design models (**Fig. 2a**). In the design model of gp100_T3 (**Fig. 2c**), there are multiple hydrogen bonds with main chains of P6 and V7, which are positioned by interactions between the intervening residues and the pMHC, and thus activation is sensitive to mutations from P4 to V7. For WT1_5, E40 of the binder makes bidentate interactions with R1, and the main chain of binder I12 makes a hydrogen bond with the side chain of peptide Y8, again consistent with the alanine scanning results. For the MART-1 target, we carried out alanine scanning experiments for two different binders (Mart-1_3 and Mart-1_43; **Fig 4.d and e**) which are centered over different regions of the target peptide; this structural shift is reflected in the alanine scanning results.

**Figure 4:**
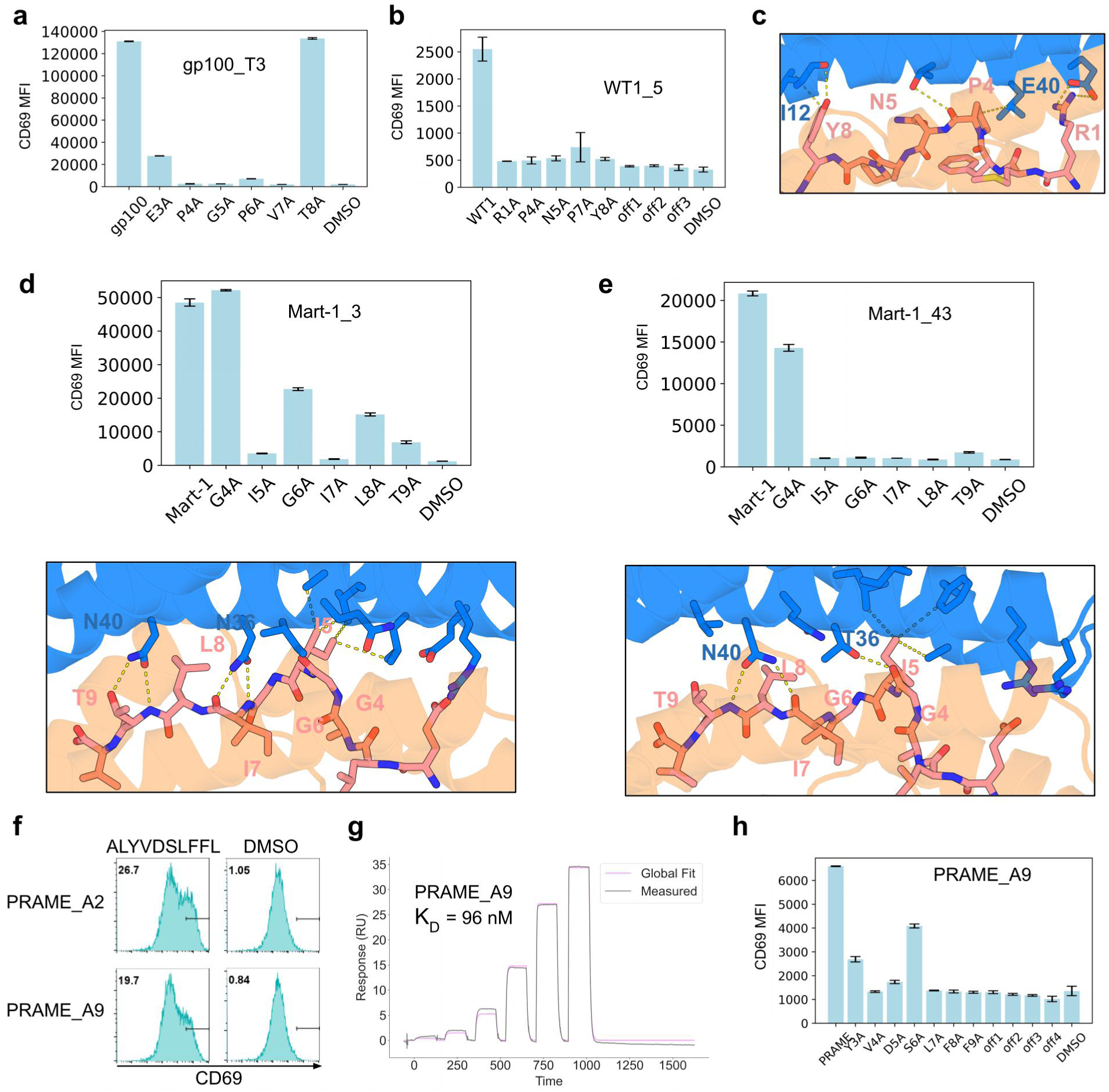
Specific activation of desTgned CARs by cognate pMHC complexes. Activation of Jurkat cells expressing the CARs by 293T cells with 5uM of different peptides measured through CD69 expression level by staining (indicated by histogram or mean fluorescence intensity (MFI}). **a,** CD69 MFI of gp100_T3 CAR with gp100 peptide, alanine mutant peptides or DMSO. **b,** CD69 MFI ofWT1_5 CAR with WT1 peptide, alanine mutant peptides, off-target peptides, or DMSO. **c,** zoom-in view of design model of WT1_5 with pMHC target. **d and e, (upper)** CD69 MFI of **d,** Mart-1_3 ore, Mart-1_43 CAR with Mart-1 peptide, alanine mutant peptides, or DMSO. **(lower)** design models of binders with Mart-1 pMHC antigen. **f,** Histograms of CD69 expression level of PRAME CARs with PRAME peptides or DMSO. **g,** SPR traces of PRAME binder A9 on PRAME pMHC monomer. **h,** CD69 MFI of Jurkat cells expressing PRAME_A9 CAR with PRAME peptide, alanine mutant peptides, four peptides with similar sequences, or DMSO.

For PRAME where we used predicted structures rather than experimental structures in the design calculations, we identified two binders (PRAME_A9, PRAME_A2) that specifically induce Jurkat activation upon PRAME peptide pulsing (**Fig.4f**). In SPR experiments, the purified designs have K_D_’s for the target pMHC of 96nM (**Fig.4g**) and 35nM (**Supp Fig.4b**). Pulsing 293T cells with the original PRAME peptide and single alanine mutant version showed that binder A9 was quite specific (**Fig.4h**). In the design model/predicted structure (**Fig.2c**), the binder interacts with the side chains of V4, S6, L7, F9 and the main chain of L7, consistent with the observed specificity.

For PRAME_A2 (**Supp Fig.4c)** and the HIV design (**Fig.2a**), one of the alanine substitutions activated more strongly than the unmutated target peptide (**Supp Fig.4d and e)**. This could reflect better loading of the mutant peptide onto MHC (*32*), or an imperfect designed interface around the residue being mutated (**Supp Fig.4c)**. Because we have structure models of all designs, the latter problem can potentially be remedied by another round of structure based design. We tested this for the PRAME_A2 design, and indeed found that activation by the target peptide over the mutant peptide could be increased by redesigning the contacting part of the design to remove a potential clash with the original peptide residue (**Supp Fig.4d).** This ability to design and optimize binders to pMHC targets without experimental structures should enable the application of our design pipeline to a very wide range of targets.

## Discussion

Given the promise of therapies targeting pMHC-I antigens, multiple approaches have been developed to identify specific binders. The central challenge is how to achieve effective targeting of the target pMHC while retaining high specificity—binding to the target peptide but not thousands of other peptides presented on the same MHC. Screening TCRs from human T cells takes advantage of the natural repertoire to ensure specificity but is fundamentally limited by availability of donors with the relevant HLAs, central tolerance which can eliminate the TCRs most activated by endogenous targets, and the overall rarity of high affinity TCRs (*1*, *5*). Engineering of existing low affinity TCRs can increase affinity but requires extra effort for controlling specificity to avoid toxicity (*26*). Screening binders from scFv libraries has primarily yielded nonspecific binders, with the most successful specific binders adopting similar docking geometry as TCRs (*1*, *3*), suggesting the importance of the docking interface for sufficient peptide contact and binding specificity. These methods have successfully identified specific binders against multiple pMHCs, but given the high cost and low hit rate they are difficult to employ against the large number of potential pMHC therapeutic targets. Our *de novo* design approach builds on lessons learned from the binding modes of TCRs and the most successful scFvs by generating binders with pMHC interaction interfaces focused on the presented peptide with limited contact with the MHC. We demonstrate that this enables robust design of specific binders for ten diverse pMHC targets; because of the simplicity of the structures (compared to scFvs and TCRs), further optimization can be carried out based on the design models/predicted structures, enabling further optimization without experimental structure determination (**Supp Fig.4c and d**). Our design pipeline is quite efficient and could be readily applicable to a wide range of pMHC targets; for example in the cases of Mart-1 and gp100, computational design took one week, and once an oligonucleotide library encoding the designs had been obtained, it took one week to identify specific binders on yeast, and two weeks to subselect specifically activating binders on Jurkat cells.

We expect the power of our design approach to rapidly design specific binders to class I pMHCs to continue to increase. First, it should be possible to learn from our design campaigns what properties correlate with specific activation on cells; for example, what types of scaffold geometries give the most effective readouts of sequence over the full peptide length. Second, as deep learning-based structure prediction, design, and model ranking methods continue to increase, it should become possible to find designs with suitable affinities and specificities in testing only a small handful of candidates. With such advances, it should become possible to generate clinical-grade pMHC binders to provide therapeutic benefits to a broad patient population, including rare mutations and HLA alleles.

## Author contributions

D.B., N.F.G., B.L. and A.M. conceived the project. D.B., N.F.G., B.L. and A.M. designed the overall experimental and computational strategies. J.B., N.F.G., B.L., and J.S. designed the binder structures and sequences. J.B., N.F.G., B.L., A.M. and J.S. performed yeast FACS assays. J.B., N.F.G., and B.L. performed mammalian cell assays, and protein purification/SPR. N.R. and D.K.V. performed yeast transformation and NGS. C.W. performed global peptide screening with MAGE-513. A.N. and G.M.V. assisted with *in silico* specificity screening using ProteinMPNN. K.C.G. assisted with project discussion and global peptide screening experimental design. D.B., J.B., N.F.G., B.L. wrote the manuscript. All authors edited the manuscript.

## Methods

### Computational design of pMHC binders

Target structures used as inputs for binder design were obtained from the Protein Data Bank for A*01:01 MAGE-A3 (PDB: 5BRZ), A*03:01 CTNNb1 (PDB: 6O9C), A*02:01 HIV-Env (PDB: 2X4O, AF3 models), A*02:01 Wilms tumor antigen 1 (PDB: 6RSY, AF3 models), A*02:01 SARS-CoV membrane protein (PDB: 3I6G), A*02:01 YFV NS4b214-22 (PDB: 6SS8), A*02:01 MART-1 (PDB: 5NHT), A*02:01 gp100 (PDB: 5EU3). The structure for A*02:01 PAP (TLMSAMTNL) was predicted by folding with a version of AlphaFold2 fine-tuned for MHC (*33*) structures; the structure for A*02:01 PRAME (ALYVDSLFFL) was predicted by folding with AlphaFold3 using all 5 model outputs. Upward-facing residues of the peptide were chosen as hotspots to condition RFdiffusion towards generating binders with high peptide contact. For scaffold recycling, a partial_t between 12 and 25 (out of 50 total RFdiffusion denoising steps) was used after docking scaffolds to new targets. ProteinMPNN was used to generate sequences for the output backbones. A range of 1,000-20,000 backbones were generated per RFdiffusion cycle, with anywhere between 4-32 MPNN sequences generated per backbone. Output structures were subject to *in silico* screening based on AlphaFold2 initial guess (pAE interaction, binder pLDDT and binder RMSD), AlphaFold2 monomer pLDDT, Rosetta contact molecular surface (CMS) per target residue or range of residues, and ProteinMPNN log probability scores of designs in complex with alanine scan mutants of the target peptide. The cutoff values varied for each target protein and round of design. This design process was iterated for each target through new rounds of partial RFdiffusion until a desired number of designs passed the cutoffs.

### AF-MHC screening

Building off previous work in predicting pMHC complexes, prior to the release of AF3, we altered the AlphaFold2 fine-tuned for MHC (*33*) to predict minibinder-pMHC complexes through additional templating of the minibinder. For screening, designs were predicted against the on-target and 2-3 relevant off-target peptides in the same HLA allele. The pAE interaction between only the peptide and the minibinder was calculated, and for each off-target a delta pAE interaction (on-target pAE – off-target pAE, more negative is better) was used to filter designs, with scores ranging from –10 to –0.5 depending on the on-target/off-target pair.

### Rosetta CMS screening

Rosetta contact molecular surface described before (*11*) uses a triangulation algorithm to calculate the contact surfaces of binder and target and gives it a score taking into account cavities and holes in the interface. For peptide contact screening, we restricted the target surface being calculated by the algorithm to the peptide only or to individual residues of the peptide, and scored its contact to the entire binder surface. Cutoff values were determined empirically for each target.

### ProteinMPNN log probability screening

ProteinMPNN was used to score the likelihood of each peptide residue being predicted when in complex with a binder. Binders were screened by scoring each peptide residue separately and filtering directly on the log probability of the on-target residue, or on the difference in log probabilities to when that residue is mutated to an alanine.

### Off-target peptide identification

To determine peptides that are likely to cross react with our peptide of interest, we used selective Cross-Reactive Antigen Presentation (*1*), and took all the outputs and cross-checked them with NetMHC4.1 (*34*), using any peptides with an EL_rank <= 0.5. Then we used a simple blossom alignment of the passing peptides against our peptide of interest to rank and choose the top off-target peptides.

### DNA library preparation

For yeast surface display, all designed protein sequences were first padded at both termini using GS to a uniform length of either 88 amino acids or 102 amino acids depending on if a 300bp or 350bp oligonucleotide library was ordered, respectively. Then the sequences were reverse translated using dnachisel while codon optimizing for *S. cerevisiae*. DNA sequences were ordered as oligonucleotide libraries via Twist Biosciences or Integrated DNA Technologies (IDT), with 300bp or 350bp sizes.

### Yeast surface display screening with FACS

Transformed S. cerevisiae EBY100 strain library cultures were grown in C-Trp-Ura (2% glucose w/v) medium and induced for expression in SGCAA (0.2% glucose w/v) medium. Cells were washed with PBSF (PBS with 1% BSA w/v) and incubated with FITC-conjugated anti-C-Myc chicken antibody (ICL, CMYC-45F) for expression sorting. For binding sorts, cells were additionally incubated with peptide-MHC biotinylated tetramer (Fred Hutchinson Cancer Center Immune Monitoring Services) or dextramer (Immudex) conjugated with phycoerythrin (PE) for on-target peptide or allophycocyanin (APC) for off-target peptide binding. When tetramers or dextramers were mixed, biotin was added to a final concentration of 1uM. Cells were incubated for 30 minutes, then washed and resuspended with PBSF. All sorts were performed on the Sony SH800 FACS instrument. FACS data was analyzed using the FlowJo software. All naive and sorted pools were sequenced using Illumina NextSeq and MiSeq and analyzed for enrichment between sorts.

### Protein binder expression and purification

Synthetic genes of top yeast display hits were ordered as eBlock gene fragments from IDT and cloned into a pET29b(+) vector containing an N-terminus His6x tag for all designs and an Avi-tag or SNAC-tag for some designs to introduce Trp to expressed sequences for protein quantification using A280 signal. Following cloning and transformation as in (*35*), we picked single colonies for sequence verification. Protein purification was similar to (*35*); briefly, we grew cultures in 50mL of Terrific Broth II with 50mg/mL of kanamycin for 6-12 hours before spiking in IPTG at 1mM final concentration and growing overnight at 18C. Cultures were harvested by spinning at >4000g for 5-10 minutes, and resuspending in lysis buffer (15mL of 25mM Tris-HCl, 300mM NaCl, 40mM Imidazole), with addition of protease inhibitor, lysozyme, and DNAse. Following lysis by sonication and centrifuge at >14,000g for 45 minutes, proteins were purified via nickel Immobilized Metal Affinity Column. Supernatant was allowed to freely drip before washing with 5mL of lysis buffer twice. Proteins were eluted with 1-2mL of elution buffer (25mM Tris-HCl, 300mM NaCl, 500mM Imidazole) before SEC using Superdex 75 10/300GL columns in HBS-EP+ buffer (0.01 M HEPES pH 7.4, 0.15 M NaCl, 3 mM EDTA, 0.005% v/v Surfactant P20, Cytiva #BR100669) and collecting relevant elution fractions.

### Surface Plasmon Resonance

Binding kinetics were analyzed via Surface Plasmon Resonance (SPR) on a Biacore 8K (Cytiva). Binding for different pMHC targets was measured by capturing biotinylated peptide-MHC monomer (Fred Hutchinson Cancer Center Immune Monitoring Services) using the Biotin CAPture Kit (Cytiva #28920234). Capture was performed by injecting 0.5 µg/mL pMHC monomer at a flow rate of 10 µL/min in HBS-EP+ (0.01 M HEPES pH 7.4, 0.15 M NaCl, 3 mM EDTA, 0.005% v/v Surfactant P20, Cytiva #BR100669) aiming for a capture level of ∼250 response units. Binder analytes in HBS-EP+ buffer were injected at a flow rate of 30 µL/min to monitor association. HBS-EP+ was also used as a running buffer during dissociation under the same flow rate conditions. Ligand concentrations ranged from 1nM to 1uM. Binding kinetics were determined by global fitting of curves to k_on_ and k_off_ assuming a 1:1 Langmuir interaction, using the Cytiva evaluation software.

### Chimeric Antigen Receptor constructs

CAR plasmids are constructed based on pSLCAR-CD19-BBZ (Addgene: 135992). In short, the configuration from N-terminal to C-terminal is CD28 signal peptide-FLAG-binder-CD8 hinge-CD28 transmembrane domain-41bb1 cytoplasmic domain-CD3Z cytoplasmic domain-P2A-mTagBFP2-P2A-PuroR.

### Mammalian cell culture and transfection

Binder sequences were reverse translated while optimizing for *H. sapiens* and ordered as CAR plasmids from Genscript based on pSLCAR-CD19-BBZ (Addgene: 135992). Lentivirus particles of the CAR plasmids were generated by transfecting 0.25mL HEK 293T cells at 800,000 cell/mL grown in DMEM medium (Gibco, 11965092) supplemented with 10% (v/v) fetal bovine serum (FBS) and 1% (v/v) penicillin–streptomycin (Pen-Strep), with 2 μl TransIT-293 (Mirus), 0.25 μg psPAX2 (Addgene, 12260), 0.1 μg pCMV-VSV-G (Addgene, 8454), and 0.4 μg plasmid in with 50 μl OptiMEM medium (Gibco, 31985070). Transfected HEK293T cells were replenished with fresh RPMI 1640 medium (Gibco, 61870036) supplemented with 10% (v/v) fetal bovine serum FBS and 1% (v/v) Pen-Strep 24 hours after transfection. Supernatant was collected 48 hours post-transfection and freeze-thawed to kill remaining HEK293T cells. To generate Jurkat-CAR cells, 0.25mL Jurkat cells (a gift from Phil Greenberg lab) at 2 million cells/mL grown in RPMI medium (10% FBS, 1% Pen-Strep) were supplemented with 8 µg/ml Protamine (Millipore Sigma, P4020) and infected with 0.4mL of collected lentivirus supernatant by centrifuging at 1,000g for 60 minutes. Lentivirus-infected Jurkat cells were replenished with fresh RPMI growth medium 24 hours post-infection and every 48 to 72 hours thereafter.

### Antigen presenting cell lines

HLA-A*01:01-expressing HEK 293T were made with lentiviral plasmid expressing human HLA-A*01:01 (a gift from Paul Thomas lab); HEK 293T constitutively expressing HLA-A*02:01 were used as HLA-A2 antigen presenting cells. HLA-A*03:01-expressing K-562 cells were made using lentiviral plasmid expression (a gift from Paul Thomas lab).

### Yeast surface display clonal binding assay

Clonal yeast surface display samples were obtained by plating sorted pools from FACS on glucose-Trp-Ura agar plates (Teknova C3260), incubating at 30C for 48 hours and picking single colonies. Cultures were grown in C-Trp-Ura (2% glucose w/v) medium and induced for expression in SGCAA (0.2% glucose w/v) medium for 12-18 hours. 50uL of culture was transferred to a 96-well plate and washed with PBSF. Cells were stained at 1:100 v/v with FITC-conjugated anti-C-Myc chicken antibody (ICL, CMYC-45F) for expression and with 1:100 v/v peptide-MHC biotinylated tetramer (Fred Hutchinson Cancer Center Immune Monitoring Services) for both on-target peptide (PE-conjugated) and off-target peptide (APC-conjugated). Cells were incubated for 30 minutes and then washed and resuspended in PBSF. All flow cytometry experiments were performed on the Attune NxT instrument and data was analyzed using the FlowJo software.

### Jurkat-CAR binding assay

50 uL of Jurkat-CAR cells at 1 million cells/mL were added to a 96-well plate. Growth medium was removed and replaced with 50 uL staining reagent made of FACS buffer (Dubecco’s PBS pH 7.2, 0.5% bovine serum albumin, and 2 mM EDTA) containing 1:100 v/v peptide-MHC biotinylated tetramer (Fred Hutchinson Cancer Center Immune Monitoring Services) for both on-target peptide (PE-conjugated) and off-target peptide (APC-conjugated). Cells were incubated for 30 minutes at 4C and then washed and resuspended in 120uL FACS buffer. All flow cytometry experiments were performed on the Attune NxT instrument and data was analyzed using the FlowJo software.

### Activation assays using flow cytometry

Antigen presenting cells (APCs) (HEK293T cells presenting HLA-A*02:01, K652 cells presenting HLA-A*03:01) were resuspended into RPMI medium at 1 million cells/mL and pulsed with peptide to final concentration 10uM (ordered from Genscript and eluted in DMSO) or equivalent volume of DMSO. 100 uL of cells were added to 96-well plates. 100 uL of Jurkat-CAR cells at 1 million cells/mL were added to MHC-presenting cells. Jurkat-CAR cells and MHC-presenting cells were incubated overnight for 12 to 18 hours. After incubation, cells were centrifuged at 800g for 2 minutes to remove supernatant. Cells were stained with 50uL reagent containing anti-mouse CD69 antibody (Biolegend, 104513) conjugated with APC in FACS buffer at 1:100 v/v for 45 minutes in 4C, then washed and resuspended in FACS buffer. CAR-Jurkat activation was measured via flow cytometry (Attune NxT) and data was analyzed using the FlowJo software.

### Global peptide scanning

A yeast display HLA-A1 library was generated as previously described (*30*) to display 9-mer peptides, with P3 and P9 as anchoring residues with limited diversity (P3 as aspartate or glutamate, P9 as tyrosine only). For other positions of the peptide library, an NNK codon was used to allow all 20 amino acids. Protein binders were expressed and purified as described with C-terminal 6XHIS tag, then biotinylated. The yeast-display HLA-A1 peptide library was selected with streptavidin-coated magnetic beads coated with biotinylated binder proteins as previously described (*30*).

## Acknowledgement

This research was supported by The Audacious Project at the Institute for Protein Design (J.B., N.F.G.), NCI Matchmakers Y1(J.B., N.F.G.), Cancer Grand Challenge from CRUK (B.L., C.W., K.C.G.), Department of the Defense, Defense Threat, Reduction Agency grant HDTRA (B.L.), Microsoft Protein Prediction (B.L.), Washington Research Foundation Fellowship (N.F.G.), NIH 2R01AI103867-11 (C.W., K.C.G.), PICI(C.W., K.C.G.), National Institutes of Health MIRA award (R35 GM142795) (G.M.V., A.N.), and the CAREER award from the National Science Foundation (grant No: 2045054) (G.M.V., A.N.). K.C.G. and D.B. are Investigators of the Howard Hughes Medical Institute. We also acknowledge Anastasia Minervina, Paul Thomas for discussion and sharing plasmids of different HLA alleles, Kevin M Jude for structural study, Colin Correnti, Phil Bradley, Long Tran, Avi Swartz, Kirsten Thompson, Linna An, Brian Coventry, Shajesh Sharma, Wei Chen, Magnus Bauer, Thomas Schlichthaerle for helping with computation and experiments. We also acknowledge Jenkins lab and Hadrup group for their coordination of co-submission.

---

**Supplemental Figure 1:**
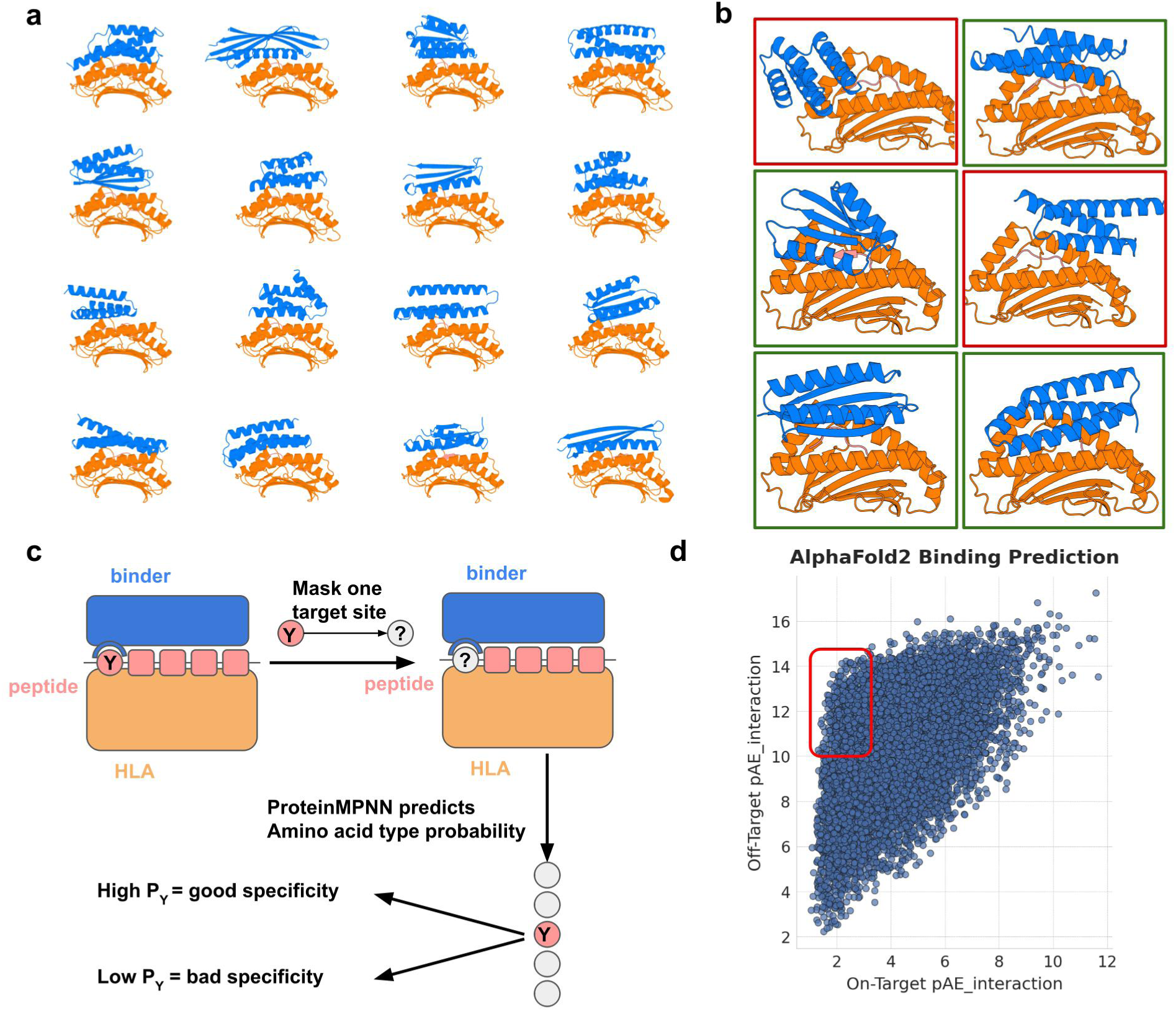
Generation of recyclable scaffold library to peptide-MHC structures. **a**, Examples of diverse diffusion scaffolds (blue) on pMHC target (orange). **b,** Designs are filtered based on peptide contact area. Designs with extensive peptide contacts (green boxes) are selected, while those with limited peptide contacts (red boxes) are filtered out. **c,** Target peptide residue mutation effect evaluation process by ProteinMPNN using predicted binder-pMHC structures. **d,** Plot of pAE_interaction values using AF-MHC to fold each design against MHC loaded with target peptides or off-target peptides. Filter (shown as red box) on low interaction pAE_interaction for on-target, high pAE_interaction for off-target.

**Supplemental Figure 2:**
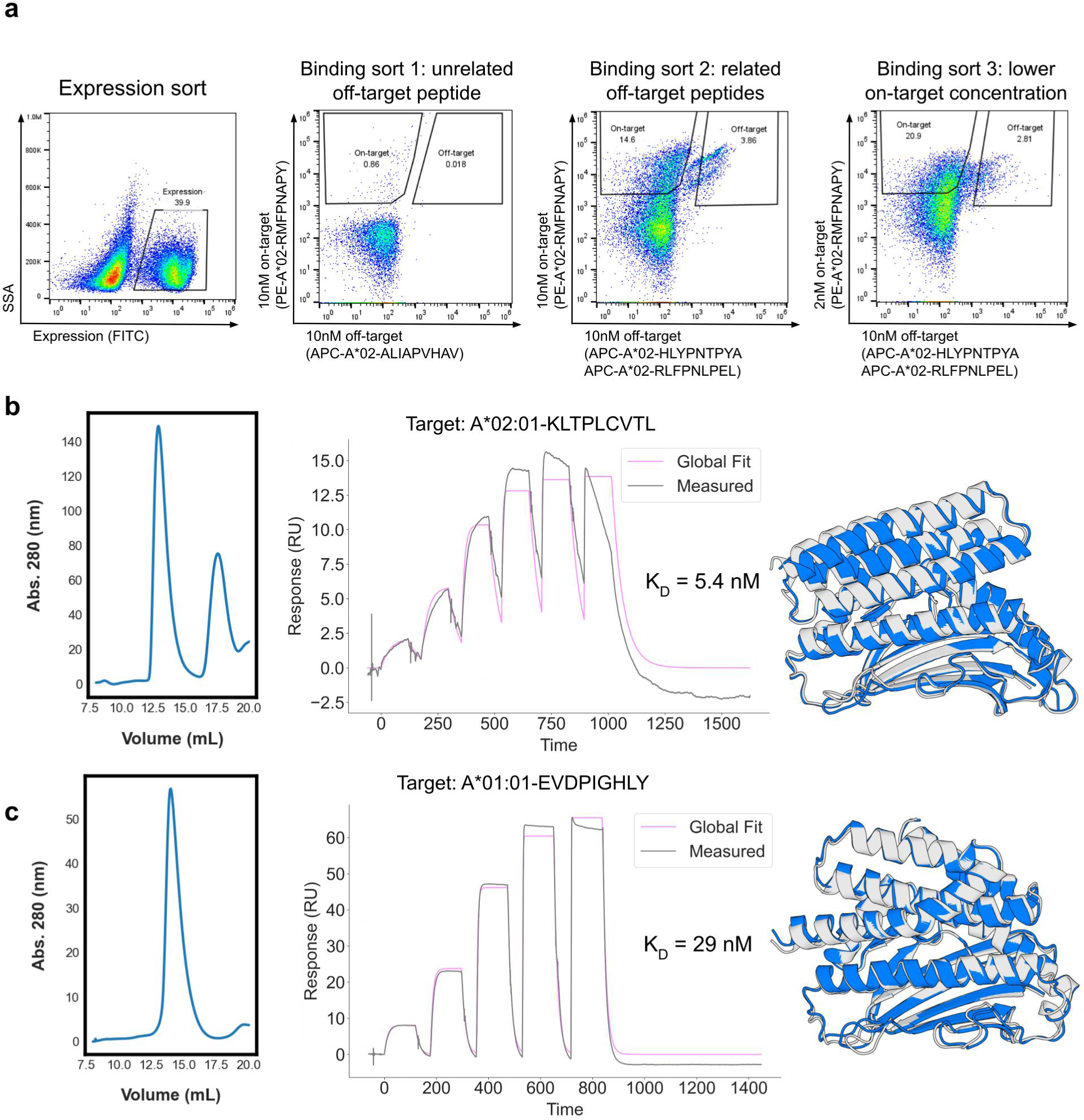
Yeast surface display library screening and biochemical characterization of individual hits. **a**, WT1 yeast surface display library screening; initial sort for surface expression followed by 3 binding sorts with target pMHC tetramer (x-axis, with decreasing concentrations for later sort) and unrelated or related off-target peptides pMHC tetramers (y-axis). Population in on-target gates was taken for subsequent sorting rounds. **b,c,** SEC trace, SPR binding kinetics (binding affinity indicated in plot), and design model (blue) overlaid with prediction (gray) for HIV-10 design against A*02:01-KLTPLCVTL **(b)** and MAGE-513 design against A*01:01-EVDPIGHLY **(c)** purified from *E. coli* expression.

**Supplemental Figure 3:**
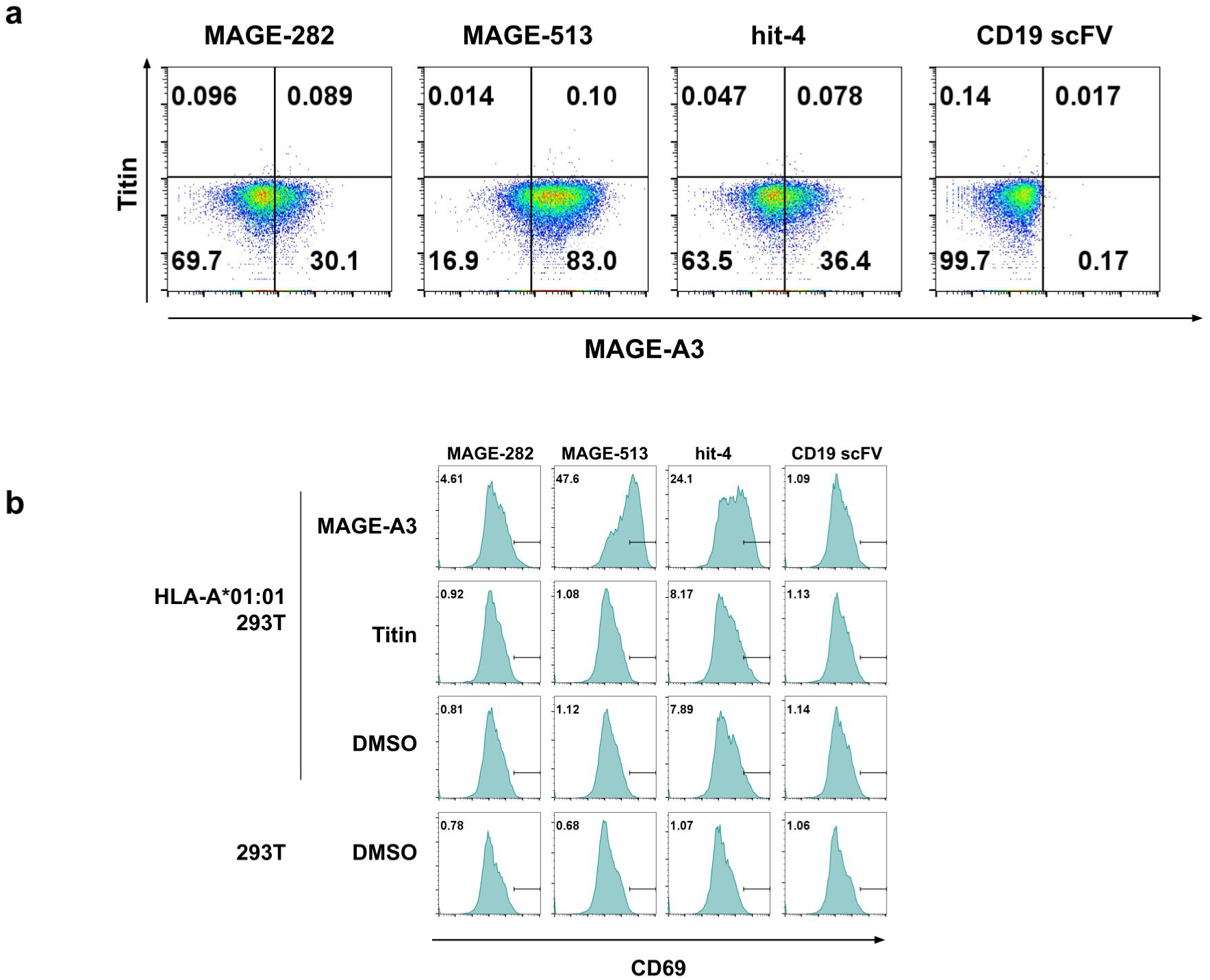
Binding and activation of different MAGE-A3 binder based CAR. **a**, Flow cytometry of jurkat cells displaying the design expressing 3 MAGE-A3 binders (MAGE-282, MAGE-513, hit-4) or CD19 scFV (negative control) based CAR incubated with on-target MAGE-A3 pMHC tetramer (x axis) and off-target Titin tetramers (y axis) at 10nM concentration. **b,** Histograms of CD69 level of Jurkat cells expressing 3 MAGE-A3 binders (MAGE-282, MAGE-513, hit-4) or CD19 scFV (negative control) based CAR incubated with 293T or HLA-A*01:01 expressing 293T upon pulsing with 5uM indicated peptides (MAGE-A3, Titin) or DMSO.

**Supplemental Figure 4:**
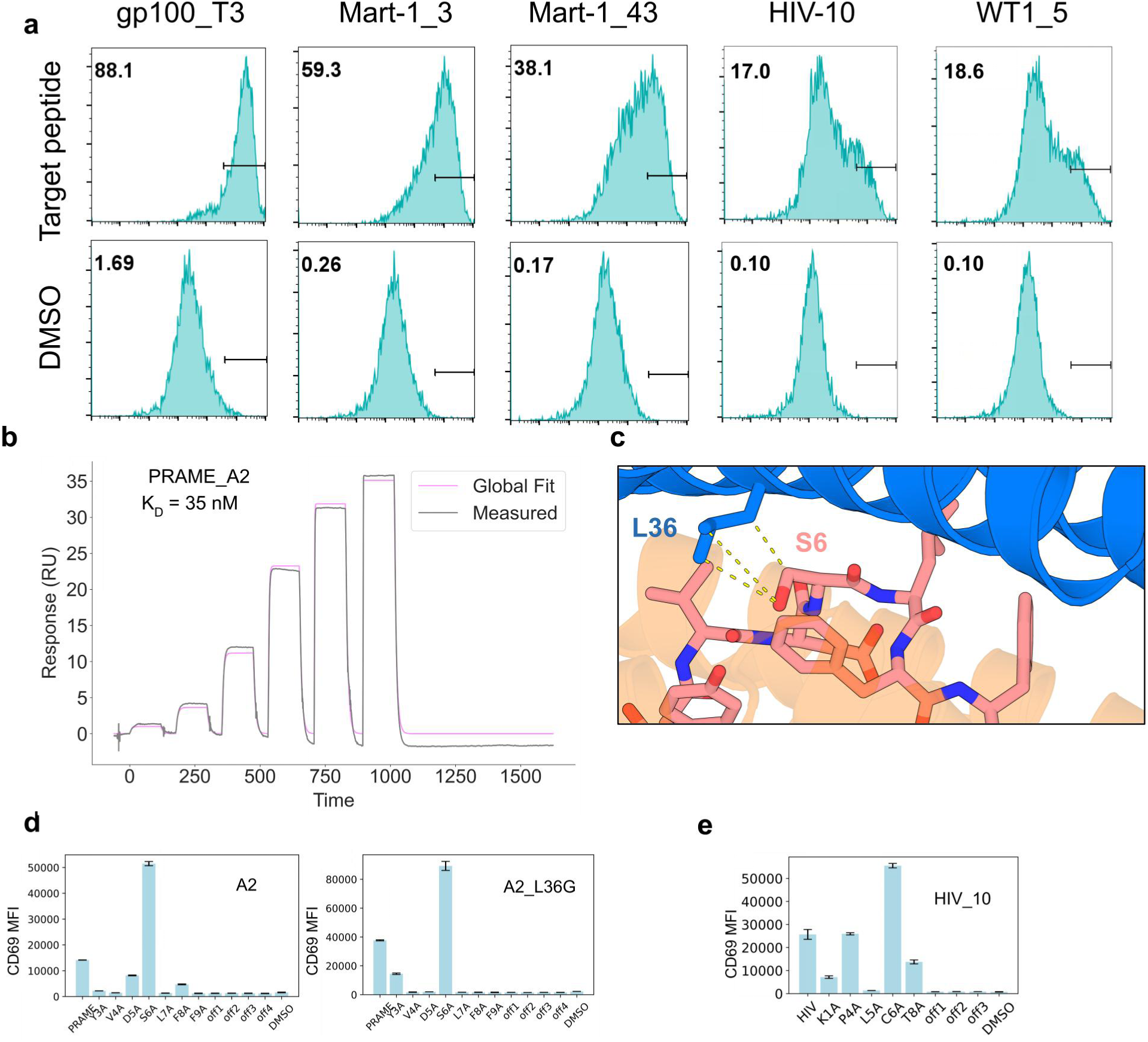
Biochemical and cellular characterization of the PRAME_A2 and HIV_10. **a**, CD69 MFI of Jurkat cells expressing indicated CARs (in each column) incubated with 293T upon pulsing with 5uM respective peptide (top) or DMSO (bottom). **b,** SPR traces of PRAME binder A2 on PRAME pMHC monomer. **c,** zoom-in view of PRAME_A2 L36 close distance with S6 of PRAME peptide form design model of PRAME_A2/PRAME peptide/HLA-A*02:01. **d,** CD69 MFI of Jurkat cells expressing PRAME binder A2 **(left)** and A2_L36G **(right)** CARs incubated with 293T upon pulsing with 5uM PRMAE peptide, alanine mutant peptides, four peptides with similar sequences, or DMSO. **e,** CD69 MFI of Jurkat cells expressing HIV binder 10 CAR incubated with 293T upon pulsing with 5uM HIV peptide, alanine mutant peptides, three peptides with similar sequences, or DMSO.

**Supplemental Table 1:**
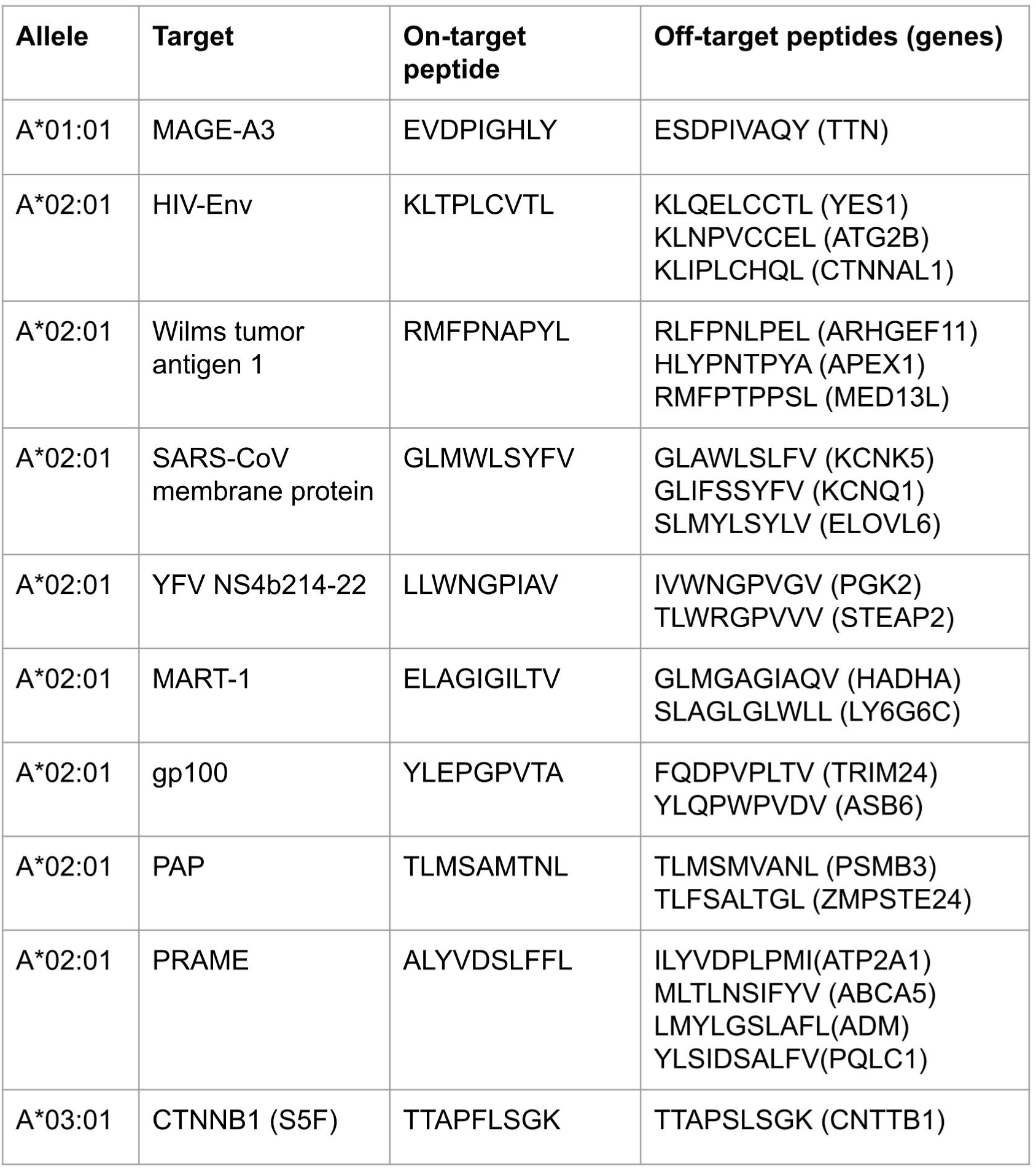
Peptide-MHC targets and selected off-target peptides. pMHC targets with respective HLA alleles, target peptides, and closely related peptides from the human proteome chosen as off-targets (gene of origin indicated in parentheses).

**Supplemental Table 2:** Design Model Characteristics. Solvent Accessible Surface Area (SASA) values in A2 for polar residues and hydrophobic residues at the interface for each targeUbinder pair, compared to TCR values when available. Delta values for SASA of MHC only and Peptide only are shown with MHC minus Peptide. RMSD of the design model to the Chai-1 or AF3 structure.

| Binder | Interface_hydrophobic_sasa (Å<sup>2</sup>) | Interface_polar_sasa (Å<sup>2</sup>) | mhc_sasa (Å<sup>2</sup>) | peptide_sasa (Å<sup>2</sup>) | Design:Prediction RMSD (Å) |
| --- | --- | --- | --- | --- | --- |
| YFV_148 | 2114 | 892 | 2692 | 777 | 0.822 |
| SARS_137 | 1516 | 716 | 1746 | 756 | 0.811 |
| HIV_10 | 1772 | 934 | 2359 | 788 | 0.933 |
| WT1_8 | 1545 | 1044 | 2248 | 761 | 0.716 |
| WT1_6rsy_tcr | 1303 | 1018 | 1960 | 769 | NA |
| PAP_116 | 1526 | 958 | 2249 | 580 | 0.454 |
| CTNNb1_15 | 1342 | 579 | 1729 | 471 | 0.463 |
| MAGE_513 | 1969 | 861 | 2488 | 793 | 0.745 |
| MAGE_5brz_tcr | 1114 | 766 | 1586 | 581 | NA |
| gp100_T3 | 1739 | 925 | 2356 | 719 | 0.63 |
| Mart1_5_5 | 1895 | 972 | 2584 | 826 | 0.754 |
| Mart1_5nht_tcr | 1231 | 626 | 1584 | 764 | NA |
| PRAME_A9 | 1998 | 807 | 2300 | 880 | 0.555 |

